# Molecular Architecture of Monkeypox Mature Virus

**DOI:** 10.1101/2024.06.21.600031

**Authors:** Ye Hong, Baoying Huang, Junxia Zhang, Cheng Peng, Weizheng Kong, Wenjie Tan, Sai Li

**Affiliations:** Beijing Frontier Research Center for Biological Structure & Tsinghua-Peking Center for Life Sciences & State Key Laboratory of Membrane Biology, School of Life Sciences, Tsinghua University, Beijing 100084, China; National Key Laboratory of Intelligent Tracking and Forecasting for Infectious Diseases, NHC Key Laboratory of Biosafety, National Institute for Viral Disease Control and Prevention, Chinese Center for Disease Control and Prevention, Beijing 102206, China

## Abstract

Since the 2022 mpox outbreak, MPXV has caused ∼100,000 infected cases with ∼200 deaths across 117 countries. and was declared a public health emergency of international concern by the WHO. The causative pathogen, monkeypox virus (MPXV), is a zoonotic double-stranded DNA orthopoxvirus. To investigate the structures and replication mechanism of MPXV, the MPXV recombinant proteins and vaccinia virus (VACV) particles have been studied by cryo-EM, however, *in situ* structural insights into major structural proteins and assembly of the authentic MPXV particles are missing. In this work, we isolated two MPXV strains from the first imported mpox case and the first local mpox case in mainland China. From the two strains, we propagated and purified mature viruses, and applied cryo-electron tomography (cryo-ET) and sub-tomogram averaging (STA) to unveil the architecture of intact MPXV virus. Interestingly, we also discovered irregularly-shaped mature virions with distorted core walls. To our knowledge, these in situ structural insights in MPXV have not been reported before.

## Main text

The mpox outbreak since 2022 has caused over 97,208 infected cases with 186 deaths across 117 countries/territories/areas and was declared a public health emergency of international concern by the World Health Organization. The causative pathogen, monkeypox virus (MPXV), is a zoonotic double-stranded DNA (dsDNA) orthopoxvirus encoding 181 proteins^1^. Currently, no specific vaccine or medicine against MPXV infection is available.

Despite efforts to elucidate the MPXV genome uncoating and replication mechanisms using recombinant proteins^2,3^, *in situ* structural insights into major structural proteins and assembly of the authentic MPXV particles are missing. Early electron microscope (EM) micrographs of the negative-stained or resin-embedded MPXV mature virus (MV), the basic form of infectious MPXV particles, have revealed its overall architecture comprising a lipid envelope coated with proteins, two lateral bodies and a biconcave viral core encapsulating genomic DNA^4^. Following fusion between MV envelope and cell membrane, lateral bodies will detach from the viral core and deliver effector proteins to host cytosol, while the viral core will be released and function as an early transcription factory protecting the virus genome^5^. Recent studies have employed vaccinia virus (VACV) for structural investigation^6-8^. However, multiple amino acid differences exist between the major structural proteins of VACV and MPXV and the currently circulating MPXV carries multiple mutations in assembly-related major structural proteins such as OPG098 and OPG136, suggesting possible variation in virus morphology^9^. Here, we optimized an MPXV propagation and purification workflow. Combining cryo-electron tomography (cryo-ET) and sub-tomogram averaging (STA), we determined the structures and assembly of palisade proteins and portal complexes to unveil the architecture of intact MPXV MVs. We also discovered irregularly-shaped MV particles with distorted core walls, which have not been reported before.

The first imported mpox case in mainland China was confirmed on September 16, 2022^10^. And the first locally acquired mpox case in the mainland China was reported in Beijing on May 31, 2023^11^. The MPXV strain MPXV-B.1-China-C-Tan-CQ01, which phylogenetically belongs to the B.1 lineage branch of IIb clade (or West African clade), was isolated from skin blister fluid of the first imported mpox case in mainland China (National Pathogen Resource Center of China preservation ID: China-CQ202209) and used for structural analysis. Briefly, virions were propagated in Vero cells in biosafety level 3 (BSL-3) laboratories, chemically fixed by paraformaldehyde (PFA), and transferred to a BSL-2 laboratory for isolation and plunge-frozen. Since the majority of MPXV MVs remain intracellular, it is necessary to lyse the infected cells before isolation to obtain sufficient virions for imaging. However, cell lysis by sonication is prohibited in most BSL-3 labs, and chemical fixation prior to cell lysing would cross-link the virions with cellular contents. As an alternative, we adjusted conventional purification method to yield MVs of high purity and concentration suitable for cryo-ET imaging (Fig. S1). The tomographic reconstructions indicated that MV particles prepared using this method remained intact and were morphologically consistent with naturally released extracellular MVs (Fig. S2).

A total of 161 MPXV MVs were imaged and reconstructed as tomographic data, revealing an oblate ellipsoidal shape of these viral particles (Table S1). Among them, 115 virions comprise a membrane protein-coated lipid envelope packaging a closed biconcave viral core and two lateral bodies, resembling the previously imaged MPXV^4^ and VACV^7,12^ MVs (Fig. 1a-d, Movie. S1). In comparison to the more rectangularly-shaped VACV MV^7^, these MPXV MVs have significantly shorter long axis and longer intermediate axis (p<0.0005), exhibiting an ellipsoidal shape with uniform dimensions (∼313×267×236 nm) (Fig. 1e). STA of palisades on viral cores revealed a 12.8-Å resolution trimer with dimensions of approximately 8×13 nm (width × height) (Fig. 1f, Fig. S3). The map fits globally well with an AF2Complex-predicted trimer of the MPXV *OPG*136-encoded residues 1-614, but exhibited local differences as described below (Fig. 1f, Fig. S4a). For simplicity, two flanks of each subunit on the top region of a palisade are named clockwise as A and B (Fig. S4b). On the top transverse plane, each two flanks (A1-B3, A2-B1, A3-B2) were connected by unassigned densities at flanks A (green arrowheads in Fig. S4b). We speculate these connecting densities are part of *OPG*130-encoded proteins, since previous investigations on VACV have confirmed multisite crosslinking interaction between OPG130 and OPG136 top region (residues 170-350), including the interaction between VACV OPG136 residues 222-233 and OPG130 residues 148-153^7,13^. Fitting the predicted MPXV OPG136 model to our map indicates that OPG136 residues 222-233 probably reside at flanks B (colored red in Fig. S4b). Meanwhile the MPXV OPG130 residues 148-153 are predicted to form an alpha helix (green arrowhead in Fig. S4c) with length similar to the unassigned density at flanks A (green arrowheads in Fig. S4b).

**Fig. 1.**
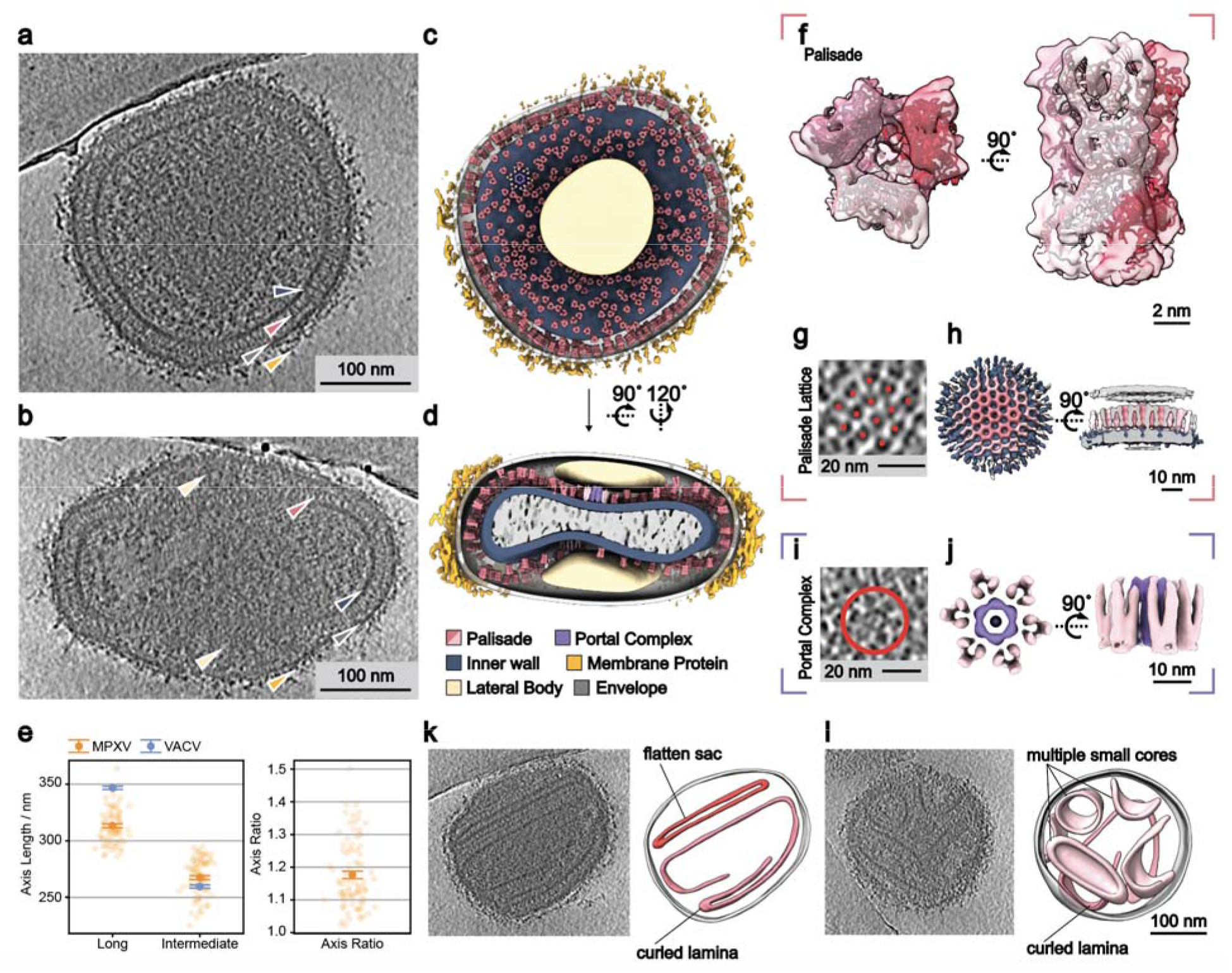
Molecular Architecture of MPXV MV. **a, b** Tomogram slices showing the top view (a) and side view (b) of two intact MV particles. Arrowheads indicate the inner wall (dark blue), palisade (pink), envelope (gray), membrane protein (orange) and lateral bodies (light yellow). **c, d** Top view (c) and side view (d) of a composite structure of the MV displayed in (a) reconstructed by projecting structures of portal complexes and palisade trimers together with envelope, membrane protein, inner wall, virion interior and lateral bodies segmented from their corresponding densities. Palisade and portal complex particles with low cross correlation value or obvious misalignment were removed. Colors corresponding to various viral components are labeled in (d). **e** Dimensional statistics of typical MPXV MV particles (n=85). Measurement of each MPXV MV is indicated by transparent orange dot. Opaque dots indicate average axis length and error bars indicate standard error of mean. Statistics of VACV from reference^7^ is plotted in blue. **f** The palisade trimer structure map fitted with an AF2Complex predicted structure of OPG136 residues 1-614. Different colors indicate three subunit of palisade trimer. **g** A tomogram slice with 5.44 nm thickness showing the honeycomb-like palisade lattice on viral core. Individual palisades are marked with red dots to show their relative arrangement. **h** Structure of the honeycomb-like lattice assembled by the palisades on viral core. The central lattice is labeled with deep pink, while the lateral palisades out of alignment mask are labeled in lighter pink. **i** A tomogram slice with 16.32 nm thickness showing the flower-shaped portal complex (marked with a red circle) identified on viral core. The tomogram was tilted to show top view of the portal complex. **j** Structure of the portal complex. The palisades are colored in light pink and the portal in purple. **k, l** Tomogram slices showing two exemplary irregularly-shaped MPXV MV particles. The envelope (gray) and core walls (pink) are segmented from the corresponding densities in the tomograms to show the irregular assembly of viral core walls. All tomograms shown here were lowpassed to 80 Å. Thickness of tomogram slices is 27.2 nm unless stated differently.

Next, we investigated the assembly of viral cores. The palisade bottom is predominantly covered with positive charges (Fig. S5a). We speculate this characteristic facilitates palisade protein’s electrostatic interaction with the underlying inner wall proteins, as reflected on the relatively high local resolution at the bottom region of palisades (Fig. S5b). Classification with larger mask indicated that 63% palisade trimers assemble into honeycomb-like lattice with an average center-to-center distance of 9 nm (Fig. 1g, h, Fig. S3), while the remaining palisades were less ordered (Fig. S3). Apart from the palisade, flower-shaped portal complexes were occasionally observed on the top and bottom surface of the MPXV viral core with no obvious distribution pattern (Fig. 1c, d, i). STA of these portal complexes revealed a density map consisting of six palisade trimers surrounding a hexagonal-cylindrical portal (Fig. 1j, Fig. S3). The average center-to-center distance of palisade proteins surrounding the portal is 10 nm (Fig. S6a), larger than that in the honeycomb-like lattice and possibly causing local distortion of palisade distribution. Lumen of the portal has an inner diameter of 6 nm on the narrowest top part and 9 nm on the widest central part (Fig. S6b). A rod-shaped density could be observed in the lumen (Fig. 1i, j, Fig. S6). To illustrate the MPXV architecture, we reconstructed a composite structure of an exemplary MPXV MV (Fig. 1c, d, Movie. S1). Notably, only particles with relatively high confidence were projected to the reconstruction, while the misaligned particles were removed.

Interestingly, the remaining 46 virions are irregularly-shaped and exhibit significantly different interior features compared to the previously reported MPXV MVs^4^ or wild-type VACV MVs^6-8^. The core walls of these particles present in forms of flatten sacs, curled laminae, or multiple small cores (Fig. 1k, l, Movie. S2, Movie. S3). To verify if the irregularly-shaped particles originated from sample preparation, we analyzed the tomograms of extracellular MVs, which were released naturally and have not been treated with detergent or sonication. In these control particles, similar features were also observed (Fig. S7a), suggesting that such features of the core wall could be natural. The current globally prevalent MPXV is the clade IIb strain, which further evolved into multiple branches. To verify whether this phenotype is specific to MPXV-B.1-China-C-Tan-CQ01, we isolated and propagated another MPXV strain of MPXV-C.1-China-C-Tan-BJ01 from the first local case (hMpxV/China/BJCDC-01/2023) in the Chinese mainland. This virus belongs to the C.1 lineage, another branch of the clade IIb strain, and is prevalent in Europe, North America, and Asia in the year of 2023^11^. This strain was propagated, purified and imaged using the same method as for MPXV-B.1-China-C-Tan-CQ01, and it also contains irregularly-shaped particles (Fig. S7b). These observations indicate a possible common feature in the assembly of MPXV during this outbreak.

In summary, here we report the molecular architecture of intact MPXV MVs. Apart from the typical MV particles, a class of irregularly-shaped MV particles packaging distorted core walls were observed in two representative MPXV strains isolated during the current outbreak. This may reflect a common feature of the currently circulating MPXV. Further investigations are needed to understand their formation mechanism and correlation to virulence. Our work broadens the understanding of orthopoxvirus assembly and provides a new aspect for orthopoxvirus morphogenesis research.

## Data and materials availability

Electron microscopy maps have been deposited in the Electron Microscopy Data Bank under accession codes EMD-XXXXX and EMD-XXXXX.

## Acknowledgements

We thank Dr. Jianlin Lei, Dr. Fan Yang and Dr. Xiaomin Li from the cryo-EM Facility, Technology Center for Protein Sciences, Tsinghua University, for their support on cryo-EM data collection. We thank the computational facility support on the cluster of Bio-Computing Platform (Tsinghua University Branch of China National Center for Protein Sciences Beijing). We thank Dr. Bingyu Liu from Imaging Core Facility, Technology Center for Protein Sciences, Tsinghua University, for her assistance on using Amira. We thank Zheyuan Zhang, Dr. Yong Chen and Dr. Yutong Song for their advices on data analysis. We thank Quanyi Wang and Yang Pan from Beijing Disease Control and Prevention, for their support on the Mpox clinical sample collection and sequence analysis. We thank our colleagues from National Institute for Viral Disease Control and Prevention, China CDC, including Wenling Wang, Li Zhao, Wen Wang, Yao Deng, Weibang Huo, Huijuan Wang Jiao Ren and Fei Ye for their support on MPXV stocks preparation in BSL-3 laboratory and Dr. Changcheng Wu for his support on sequence analysis of MPXV.

## Author Contributions

S.L. and W.T. conceived and supervised the project. B.H. isolated and propagated the virus. Y.H. and J.Z. purified the virions, prepared the cryo-sample and collected the EM data. Y.H. and C.P. analyzed the structures and statistics. Y.H., B.H., J.Z., W.K. and S.L. wrote the manuscript. All authors critically revised the manuscript.

