## Supplementary Info for "Molecular Architecture of Monkeypox Mature Virus"

**Materials and Methods**

**MPXV Isolation and Propagation**

MPXV isolation and propagation were performed in biosafety level 3 (BSL-3) laboratories of the Viral Disease Control and Prevention, China CDC. The MPXV strain MPXV-B.1-China-C-Tan-CQ01 was isolated from the first imported mpox case in mainland China^1^. The MPXV strain MPXV-C.1-China-C-Tan-BJ01 was isolated from the first local mpox case in mainland China^2^. Specimens were diluted at a 1:5 ratio in Minimum Essential Medium (MEM) supplemented with 2% fetal bovine serum and 1% antibiotics (penicillin 5,000 IU/mL, streptomycin 2,500 μg/mL, and amphotericin B 10 μg/mL, collectively referred to as PSA), and maintained at 25℃ for 30 min before being inoculated into the Vero cells. After inoculation, the cells were incubated with the diluted specimens for 6 hours at 37℃ in 5% CO_2_, and subsequently cultured for 96 hours in refreshed medium. The isolated virus was identified through cytopathic effect observation, quantitative polymerase chain reaction and immunofluorescence assay. Plaque assays were utilized to quantify infectious plaque-forming units (PFUs).

For MPXV propagation, Vero cells were cultured in T-175 flasks and infected with a multiplicity of infection of 0.01, then cultivated in MEM containing 2% fetal bovine serum for four days at 37 ℃ in 5% CO_2_. MPXV cultures were harvested once all cells exhibited cytopathic effect without significant detachment or flotation. Virus titers were determined by plaque assay, ranging between 2×10^6^ and 1×10^7^ PFU/ml.

**MPXV Sample Preparation**

To obtain sufficient MPXV MVs for cryo-ET imaging, infected cells were suspended in phosphate-buffered saline (PBS) and subjected to three freeze-thaw cycles (-80 ℃ to 37 ℃) to facilitate the release of intracellular virus particles. Subsequently, cell debris was removed by centrifugation at 4 ℃ and 2,000 × g for 10 minutes, and the released virus particles in the supernatant were fixed with 3% PFA at 4 ℃ for 48 hours within the BSL-3 laboratory.

Upon complete inactivation, the virions were transferred to a biosafety level 2 (BSL-2) laboratory for isolation. First, the virions were pelleted through a 36% sucrose cushion by ultracentrifugation (Beckman, IN) at 4 ℃ and 32,900 × g for 100 minutes. Next, the pellet was treated with 1% Triton X-100 at 37 ℃ for 2 h to alleviate inter-virion crosslinking, and concentrated by ultracentrifugation through 36% sucrose cushion. Subsequently, the sample was treated with a no-touch ultrasonic homogenizer (SCIENTZ, CN) at 80% power in core buffer (10 mM Tris-HCl pH 9.0, 1 mM DTT). The separated virus particles were then purified by 15%-45% sucrose-density gradient ultracentrifugation at 26,000 × g for 60 min and fractionated with a TRIAX flow cell (BioComp, CA). The band containing virions was collected, further treated with 1% NP-40 at 37 °C overnight, concentrated through ultracentrifugation at 32,900 × g for 60 min and finally resuspended in distilled water. The main procedure was shown in Fig. S1, method A.

Extracellular MVs that were naturally released were isolated directly from the infected cell supernatant. For MPXV-B.1-China-C-Tan-CQ01 strain, the sample was purified by ultracentrifugation first through 36% sucrose cushion, and then 15%-45% sucrose-density gradient. The virus fraction was diluted in PBS, sedimented at 32,900 × g for 60 min to remove sucrose and finally resuspended in distilled water (Fig. S1, method B). For MPXV-C.1-China-C-Tan-BJ01 strain, purification followed the same steps, except that density gradient centrifugation was not performed.

**Cryo-electron Tomography**

For cryo-ET sample preparation, purified virions were sonicated for 3 min and centrifuged at 5,000 rpm for 1 min to remove large aggregates. A glow-discharged 200 mesh holey carbon film coated copper grid (R2/2; Quantifoil, Jena, Germany) was soaked into 10 µL purified virus and incubated at room temperature for 60 min. Then, 4 µL distilled water was supplemented. The grid was blotted for 3 s and plunge frozen in liquid ethane using a Cryo-plunger 3 (Gatan, CA).

The grids were imaged using a 300 kV Titan Krios electron microscope (Thermo Fisher Scientific, Hillsboro, OR) equipped with GIF Quantum energy filter (slit width 20 eV) and K3 direct electron detector (Gatan, CA). Data for sub-tomogram averaging (STA) was collected under 64,000× magnification and super resolution mode, resulting in a calibrated pixel size of 1.36 Å. Tilt series were collected using the dose-symmetric scheme in SerialEM^3^ with a tilt range from -51° to 51° and a step size of 3°. For each tilt, a movie consisting of 10 frames was recorded at an exposure of 3.94 e^-^/Å^2^, resulting in a total dose of 137.9 e^-^/Å^2^ per tilt series.

**Cryo-electron Tomography Data Processing and Sub-tomogram Averaging**

Tilt series of the detergent-treated MPXV-B.1-China-C-Tan-CQ01 MVs were collected and were preprocessed using an in-house developed software. In brief, beam-induced motion was corrected by averaging the last 9 frames of each movie using MotionCor2^4^. Defoci of the averaged movies were estimated using CTFFIND4^5^. Tilt series were then aligned using AreTomo 1.2.5^6^, three-dimensional contrast transfer function (CTF) corrected and reconstructed using NovaCTF^7^.

The viral core walls of typical MVs from 105 tomograms were manually segmented in Dynamo^8^. For STA, 21,266 particles were oversampled along the segmented palisade layer. Additionally, 3,339 top-view particles were manually selected in IMOD^9^. Initial alignment of the palisade lattice was performed in Dynamo^8^ without symmetry imposed under 4-fold binning. After observing a clear C6-like palisade arrangement, C3 symmetry was applied for further alignment, resulting in a global STA density map illustrating a honeycomb-like palisade lattice structure.

To distinguish honeycomb-like lattice from less ordered palisade trimers, all particles underwent random in-plane rotation before being classified in Dynamo with two references created as described below. The honeycomb-like palisade lattice obtained above was C360 symmetrized and low-pass filtered to serve as a reference for the less ordered palisades. The central palisade trimer of the C360 symmetrized map was extracted as a single unit, and seven units were aligned to the honeycomb-like lattice to establish a reference for the honeycomb-like lattice structure.

For STA of single palisade trimer, the central palisade trimer of the above aligned lattice was masked and randomly rotated in-plane for further alignment with C3 symmetry imposed. After obtaining a solid density map, we cropped the single palisade trimer under 2-fold binning for finer alignment. The final map was reconstructed and sharpened with a b-factor of -1,766.47 Å^2^ in Relion 4.0^10^. The map displayed in the figure was dust-hided and the mask used for sharpening was applied to it. After each oversampling project, geometric restriction was applied to remove outlying or misoriented particles.

For STA of the portal complex, 42 top-view portal complexes were manually annotated in IMOD^9^ for initial alignment in Dynamo^8^, resulting in a preliminary map resembling a previously reported VACV portal complex structure (emd_18917)^11^. To obtain side-view particles from oversampled particles, two independent multireference alignment projects were done to distinguish portal complex from palisade proteins. The first project classified portal complex particles from the palisade lattice under 4-fold binning, based on the reference of initially aligned top-view portal complex from manually picked particles. The second project classified portal complex particles from single palisade trimers under 2-fold binning, based on the reference of emd_18917. The particles classified as portal complexes in both classifications underwent further analysis. Those with a coordinate difference less than 10.88 nm and an orientation difference smaller than 30° between two alignments were extracted and combined with the manually picked particles for final alignment and average. The map displayed in the figure was segmented and its surrounding noise was masked to display with ChimeraX^12^. The summary of STA procedure is shown in Fig. S3.

**Density Map Segmentation and Whole Virus Projection**

Virus envelope, lateral bodies, inner wall and its interior densities were segmented manually from IsoNet^13^-processed tomograms using 3D Slicer^14^. Membrane proteins were identified automatically with ilastik^15^ and density contacting virus envelope was selected for display with Amira^16^. Density maps displayed in figures were generated in ChimeraX^12^. Segmentation within density maps shown in figure 1h and 1j was done in ChimeraX^12^. Projection was done in Chimera^17^ using an in-house Matlab script. Palisade trimers selected for 2-fold binning alignment and manually picked portal complexes were projected and displayed.

**AF2Complex Prediction**

Atomic model used in fitting was predicted by AF2Complex 1.4^18^. Residues 1-614 of MPXV-B.1-China-C-Tan-CQ01 OPG136 was predicted as a trimer, and residues 1-281 of MPXV-B.1-China-C-Tan-CQ01 OPG130 was predicted as a monomer.

**Quantification and Statistical Analysis**

The long axis and intermediate axis of MPXV MVs were measured in IMOD^9^. MPXV dimensions were compared with VACV dimensions via Weltch’s t-test. All VACV dimensions were analyzed based on the mean value, standard error of mean and event number of VACV measurements previously reported by M. Hernandez-Gonzalez. *et al.*^11^ Comparisons between dimensions of MPXV MVs purified via two methods shown in Figure S2b were performed in Python.

Statistical plots were generated in Python unless stated otherwise.

**Supplementary Figures**

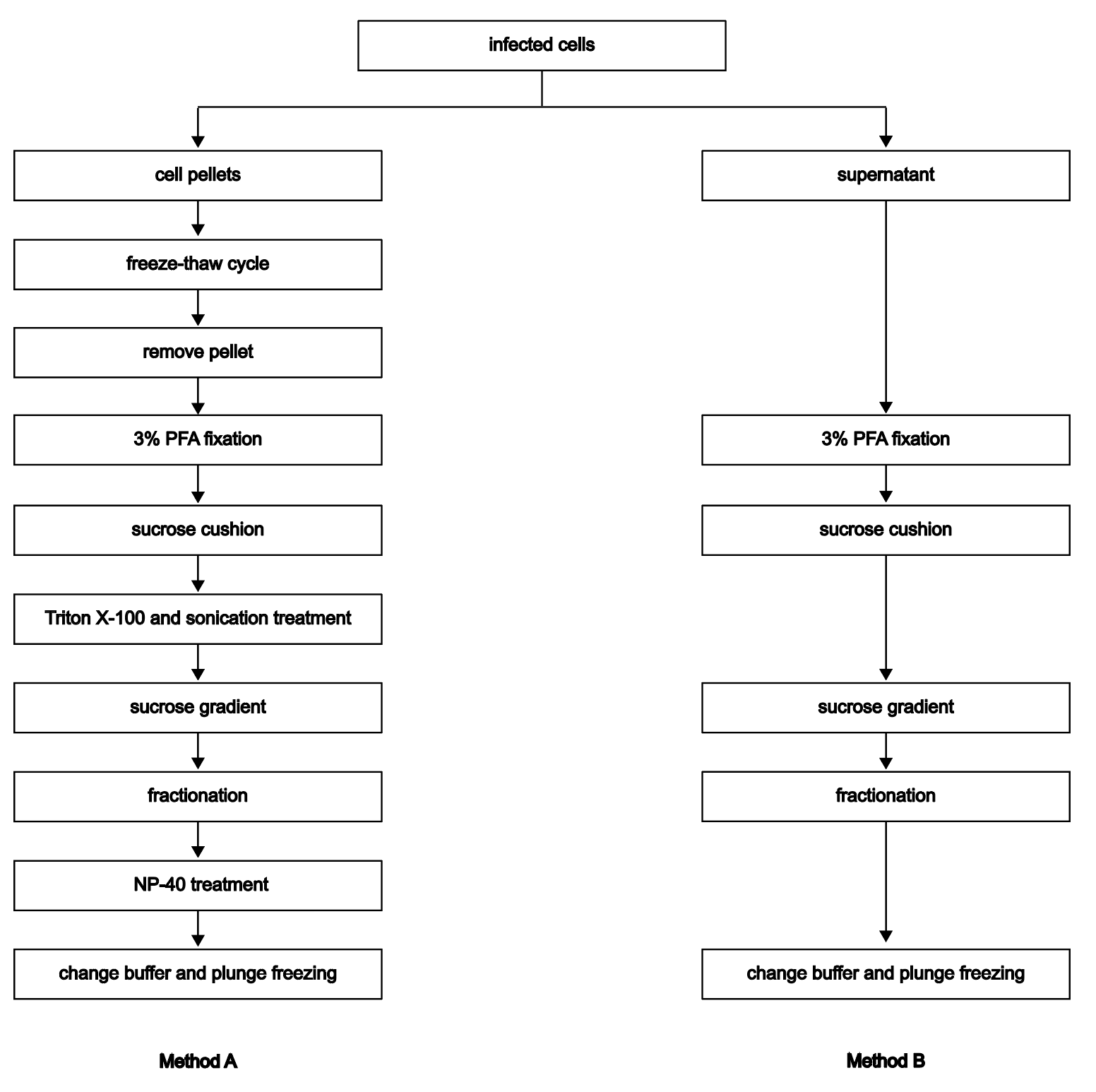

**Fig. S1** Purification workflow of MPXV-B.1-China-C-Tan-CQ01 strain MV particles.

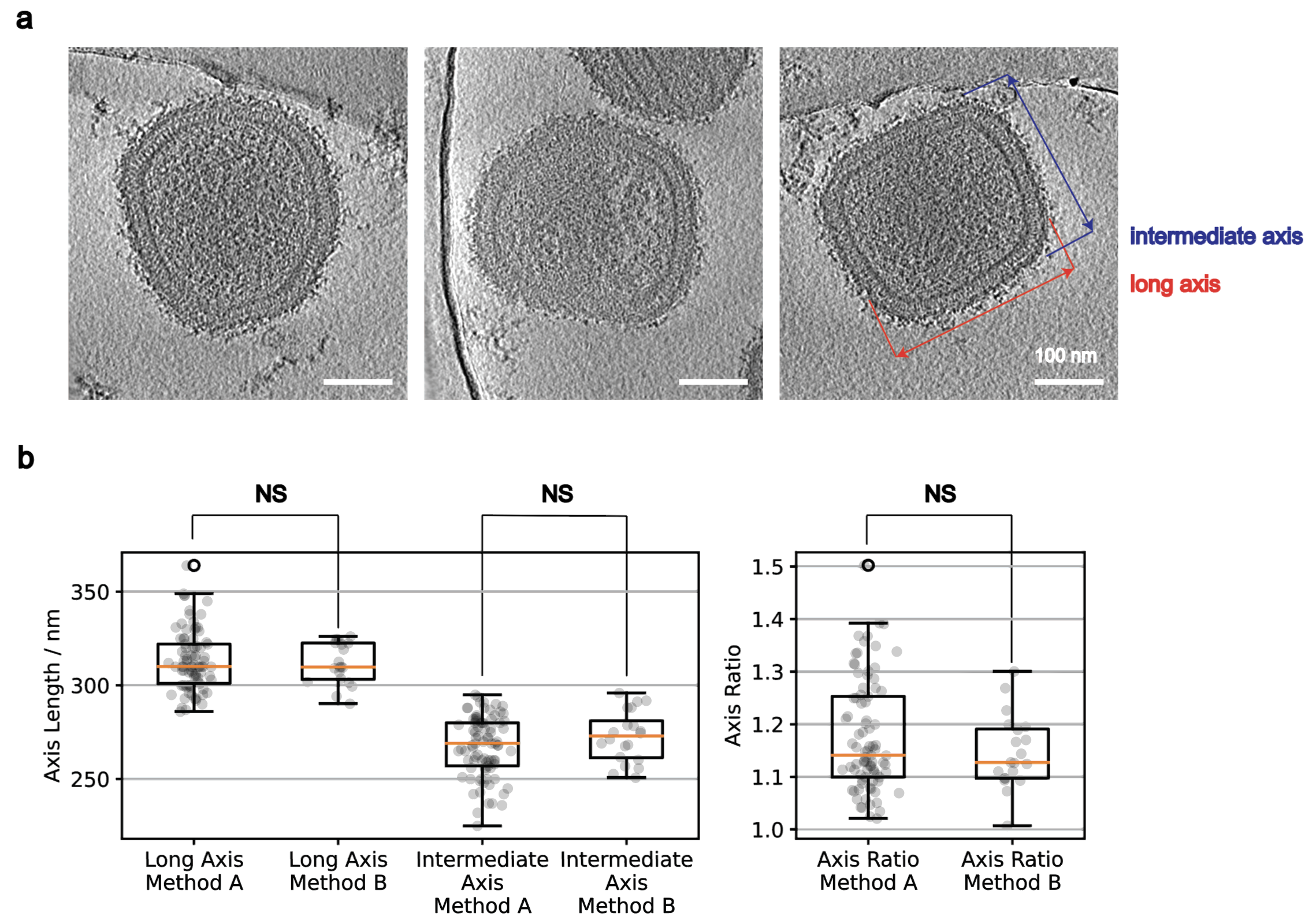

**Fig. S2** MPXV purified via two methods is similar in shape and size. **a** Extracellular MVs naturally released were purified without detergent treatment and sonication via Method B in Fig. S1. Thickness of tomogram slices is 27.2 nm. **b** Comparison between particle dimensions and axis ratio of MPXV MVs purified via two methods. Two sample independent t-test was done between measurements of 85 MVs purified via Method A and 20 MVs purified via Method B. NS represents the difference between two samples is not significant (two-sample independent t-test p-value>0.05).

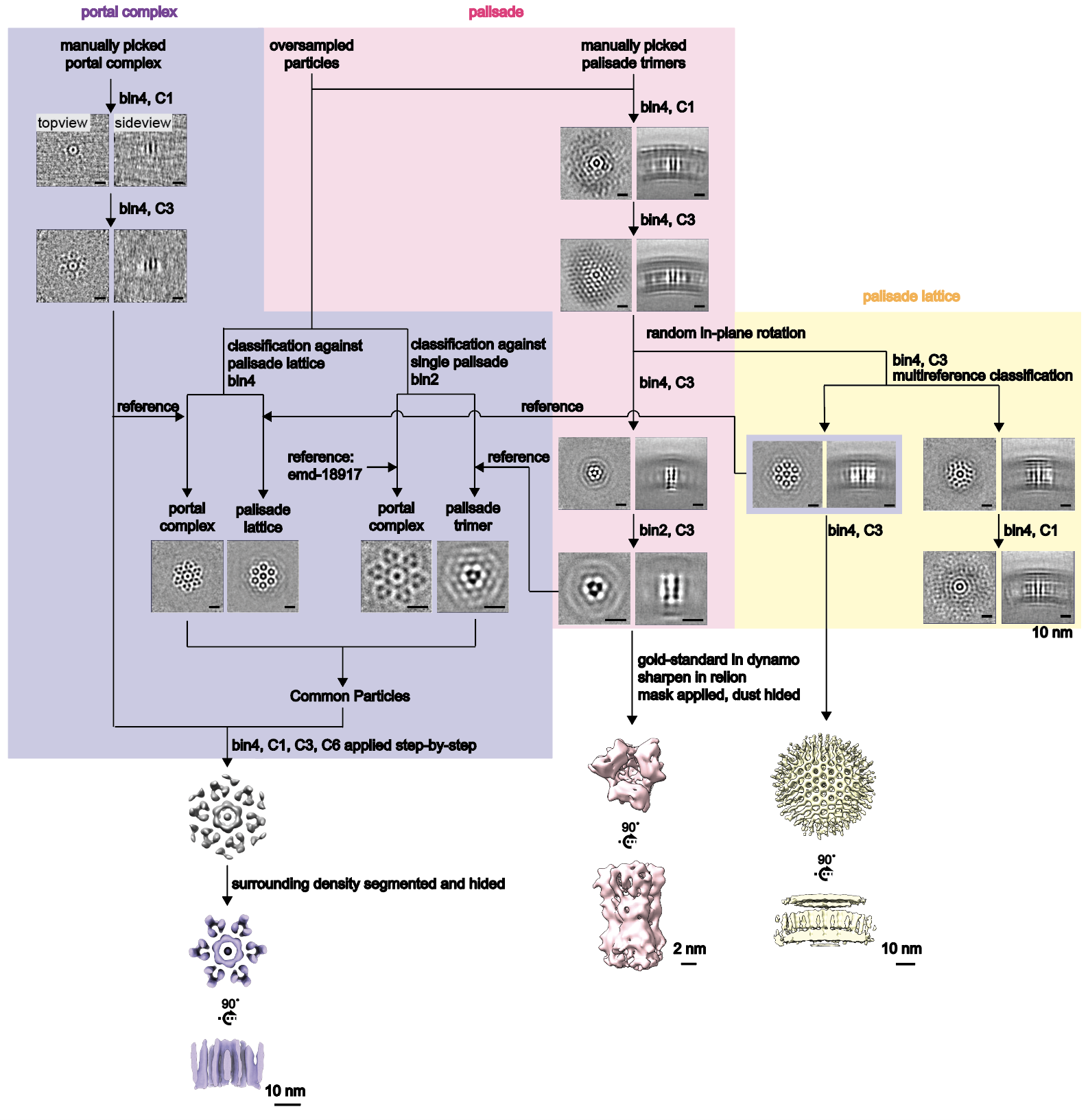

**Fig. S3** Summary of sub-tomogram averaging (STA) process.

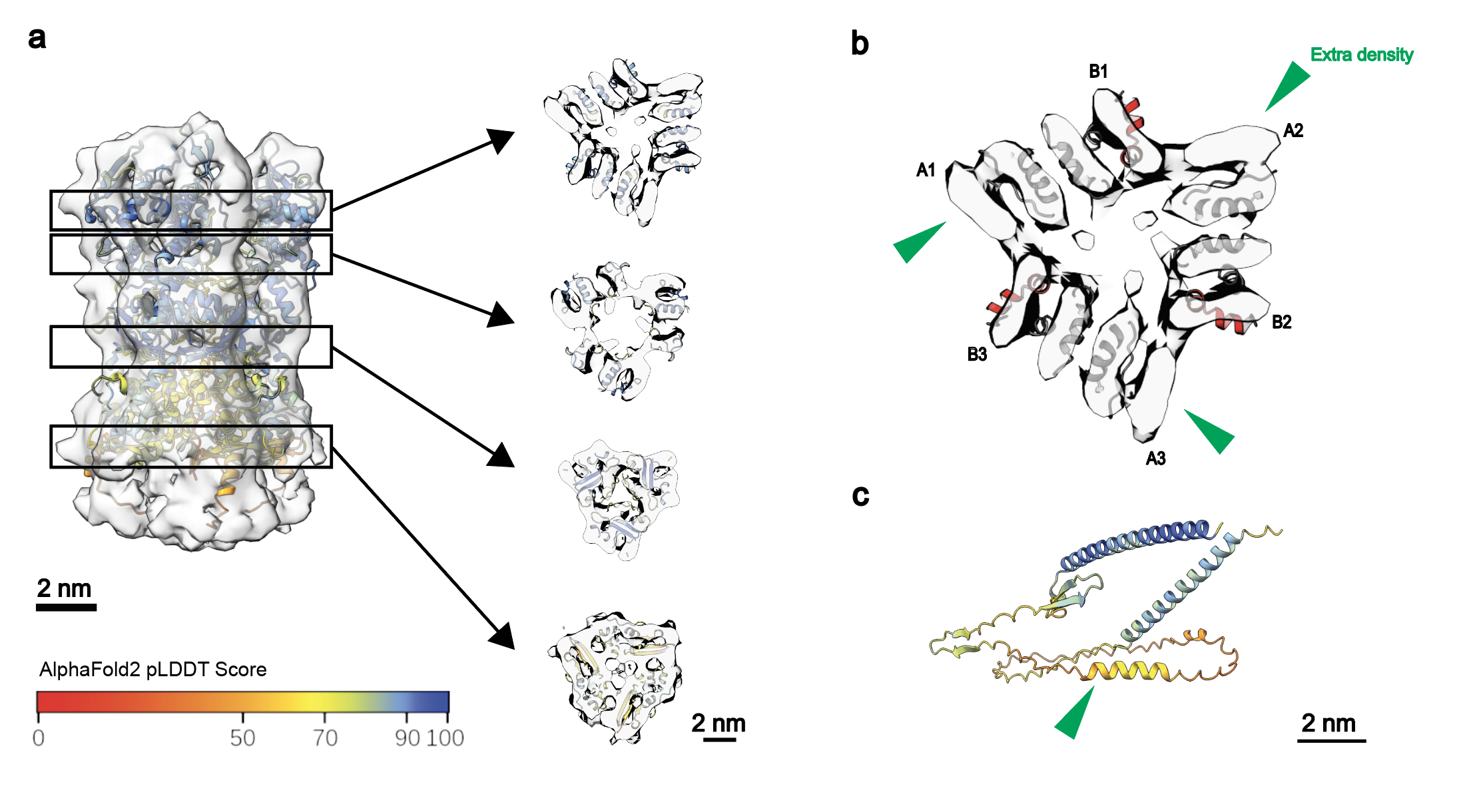

**Fig. S4 a** STA density map and fitted OPG136 model were sliced at four different transverse planes and zoomed-in for display. The predicted model is colored according to pLDDT score. **b** Magnified top slice in **a**. Residues 222-233 were colored in red. Extra densities which could not be assigned to the predicted model were indicated with green arrowheads. A1, A2, A3 and B1, B2, B3 represent flank A and B of each three subunits in a palisade trimer. **c** AF2Complex prediction of OPG130 structure. Green arrowhead indicates residues 148-153. The predicted model is colored according to pLDDT score.

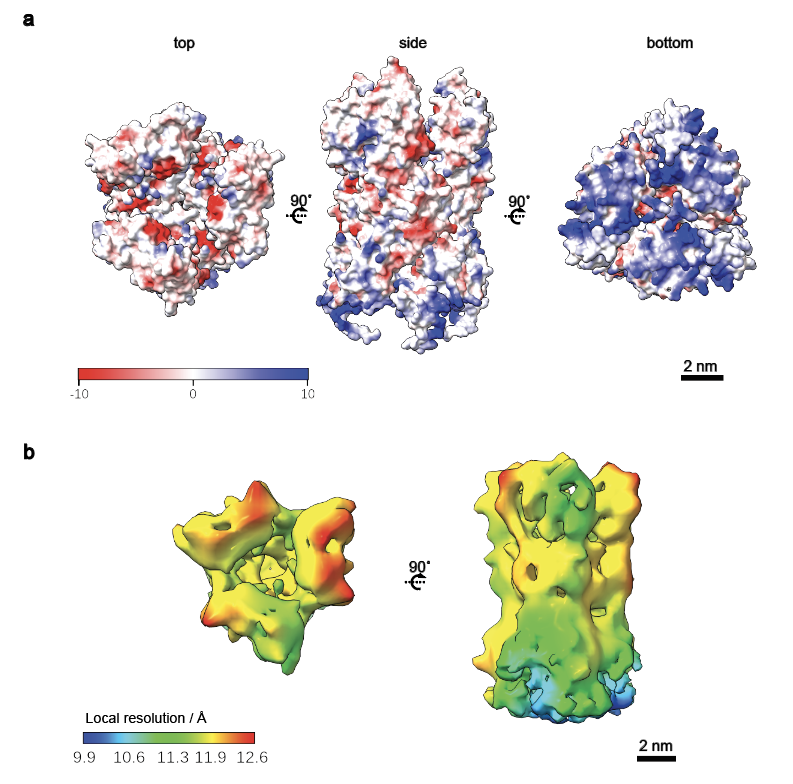

**Fig. S5 a** Surface charge of predicted palisade trimer. **b** Local resolution of palisade trimer STA density map.

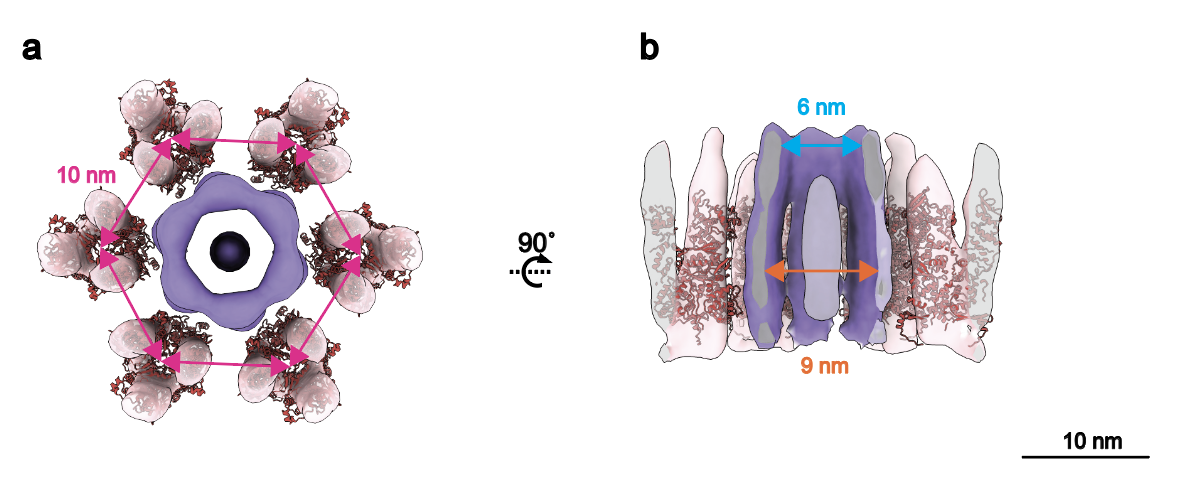

**Fig. S6** Portal Complex fitted with surrounding palisade trimers. **a** Top view of the portal complex STA structure. Average distance between palisade trimers in portal complex is 10 nm. **b** Sliced side view of the portal complex STA structure. The narrowest and widest inner diameters of the portal lumen are 6 nm and 9 nm respectively.

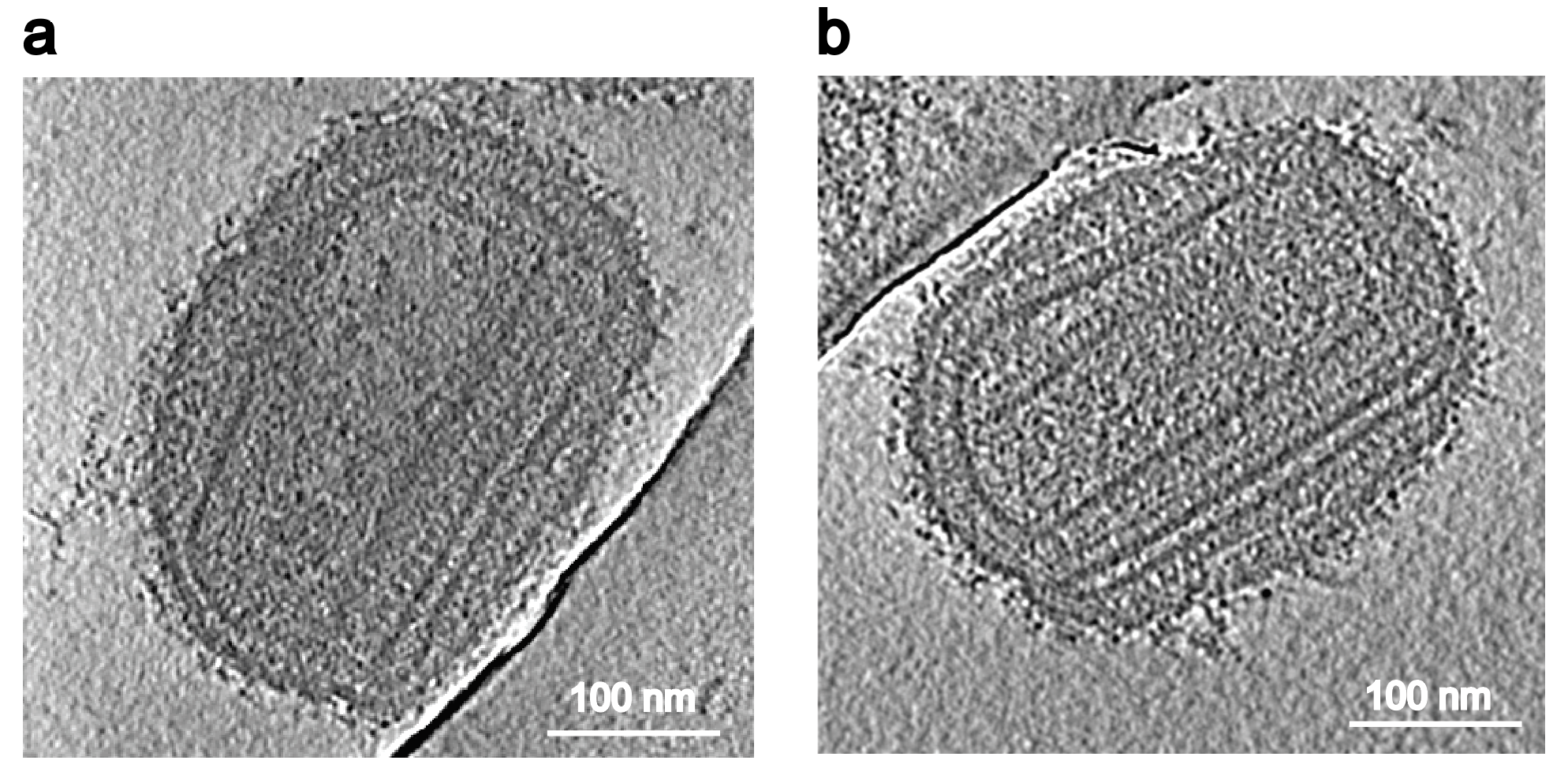

**Fig. S7** Irregularly-shaped MV particles from extracellular MPXV that were released naturally. **a** A tomogram slice of MPXV-B.1-China-C-Tan-CQ01 particle isolated without detergent treatment and sonication (Fig S1, method B). Tomogram slice thickness is 27.2 nm. **b** A tomogram slice of MPXV-C.1-China-C-Tan-BJ01 particle (without detergent treatment, sonication or sucrose-gradient ultracentrifugation). Tomogram slice thickness is 19.7 nm.

**Movie. S1** An examplary virion and its reconstructed composite structure are shown in movie. Tomogram was lowpassed to 80 Å and virus components were labeled.

**Movie. S2** An examplary irregularly-shaped particle with flatten sac and curled lamina is shown in movie. Tomogram was proccessed by isonet and the distorted core walls inside were segmented and labeld.

**Movie. S3** An examplary irregularly-shaped particle with multiple small cores and curled lamina is shown in movie. Tomogram was proccessed by isonet and the distorted core walls inside were segmented and labeld.

**Table. S1** Cryo-ET data collection and reconstruction statistics.

| **Data collection** | | | | | | | |
| --- | --- | --- | --- | --- | --- | --- | --- |
| **Microscope** | Titan Krios | | **Detector** | | Gatan K3 | | |
| **Magnification** | 64,000 | | **Pixel size (Å)** | | 0.68 (super-resolution) | | |
| **Voltage (kV)** | 300 | | **Energy filter** | | Gatan GIF Quantum, 20 eV slit | | |
| **Tilt Range** | -51° ~ 51° | | **Number of tilts** | | 35 | | |
| **Frames per tilt** | 10 | | **Tilt schemes** | | Dose-symmetric | | |
| **Exposure (e-/Å2)** | 137.9 | | **Defocus range (μm)** | | -2.0 ~ -5.5 | | |
| **Software** | SerialEM | | | | | | |
| **Reconstruction and Sub-tomogram Averaging** | | | | | | | |
| **STA** | | Dynamo 1.1.333, Relion 4.0 | | | | | |
| **Number of tomograms** | | 210 | | | | | |
| **Number of typical virions** | | 115 | | | | | |
| **Dataset** | | Palisade  trimer | | Honeycomb-like lattice | | Less ordered palisade | Portal complex |
| **Final number of particles** | | 16,074 | | 10,502 | | 6,073 | 623 |
| **Symmetry imposed** | | C3 | | C3 | | C1 | C6 |
| **Resolution (Å)** | | 12.8 | | N/A | | N/A | N/A |
| **B-factor (Å2)** | | -1,766.47 | | N/A | | N/A | N/A |
| **Final pixel size (Å)** | | 2.72 | | 5.44 | | 5.44 | 5.44 |
| **Number of Irregularly-shaped virions** | | 46 | | | | | |
